# Gene composition as a potential barrier to large recombinations in the bacterial pathogen *Klebsiella pneumoniae*

**DOI:** 10.1101/255349

**Authors:** Francesco Comandatore, Davide Sassera, Sion C. Bayliss, Erika Scaltriti, Stefano Gaiarsa, Xiaoli Cao, Ana Gales, Ryoichi Saito, Stefano Pongolini, Sylvain Brisse, Edward Feil, Claudio Bandi

## Abstract

*Klebsiella pneumoniae* (Kp) is one of the most important nosocomial pathogens world-wide, being responsible for frequent hospital outbreaks and causing sepsis and multi-organ infections with a high mortality rate and frequent hospital outbreaks. The most prevalent and widely disseminated lineage of *K. pneumoniae* is clonal group 258 (CG258), which includes the highly resistant “high-risk” genotypes ST258 and ST11. Recent studies revealed that very large recombination events have occurred during the recent emergence of Kp lineages. A striking example is provided by ST258, which has undergone a recombination event that replaced over 1 Mb of the genome with DNA from an unrelated Kp donor. Although several examples of this phenomenon have been documented in Kp and other bacterial species, the significance of these very large recombination events for the emergence of either hyper-virulent or resistant clones remains unclear. Here we present an analysis of 834 Kp genomes that provides data on the frequency of these very large recombination events (defined as those involving >100Kb), their distribution within the genome, and the dynamics of gene flow within the Kp population. We note that very large recombination events occur frequently, and in multiple lineages, and that the majority of recombinational exchanges are clustered within two overlapping genomic regions, which result to be involved by recombination events with different frequencies. Our results also indicate that certain non-CG258 lineages are more likely to act as donors to CG258 recipients than others. Furthermore, comparison of gene content in CG258 and non-CG258 strains agrees with this pattern, suggesting that the success of a large recombination depends on gene composition in the exchanged genomic portion.

**Author Summary:** *Klebsiella pneumoniae* (Kp) is an opportunistic bacterial pathogen, a major cause of deadly infections and outbreaks in hospitals worldwide. This bacterium is able to exchange large genomic portions (up to a fourth of the entire genome) within a single recombination event. Indeed, the most epidemiologically important Kp clone, is actually a hybrid which emerged after a > 1Mb recombination event. In this work, we investigated how recombinations affected the evolution of the most studied Kp Clonal Group, CG258. We found that large recombinations occurred frequently during Kp evolution, and occurred preferentially in a well-delimited genomic region. Furthermore, we found that four epidemiologically important clones emerged after large recombinations. We identified the donors of several large recombinations: despite many Kp lineages acted as donors during CG258 evolution, two of them have been involved more frequently. We hypothesize that the observed pattern of donors-recipients in recombinations, and the presence of a large recombinogenic region in Kp genome, could be related to gene composition. Indeed, genomic analyses showed a pattern compatible with this hypothesis, suggesting that gene content can represent a main factor in the success of a large recombination.

## Introduction

Recent genomic studies revealed that some bacterial species are capable of exchanging large (> 5 Kb) and/or very large (>100Kb, up to over 1 Mb) genomic portions [1] provoking sudden and extended changes into the recipient genome, e.g. the acquisition and/or loss of several genes and multiple nucleotide variations [2]. In particular, genomic studies published in 2014 and 2015 described large and very large recombination events (of up to 20% of the entire genome) involving epidemiologically relevant strains of *Klebsiella pneumoniae* (Kp), an important nosocomial pathogen [3–7].

Kp is a Gram-negative bacterium and member of the *Enterobacteriaceae* family. This species has a diverse ecology: in addition to being a common colonizer of the guts of humans and other mammals (including livestock), it is also associated with invertebrates, plants, and multiple niches in the environment including soil and water [8, 9]. Kp is comprised of three major phylogroups, named KpI, KpII and KpIII [10]. Although KpII and KpIII have been defined as species *K. quasipneumoniae* and *K. variicola* [11, 12], for simplicity we will still refer to them as *K. pneumoniae* groups II and III. Analysis of a diverse global dataset revealed ecological differences between these groups; while KpI strains are often isolated from hospitalized human patients, KpII strains are frequently associated with healthy carriers, and KpIII strains are mainly found in the environment or in association with other mammals or plants [9]. Kp can behave as an opportunistic pathogen, especially in immuno-compromised hosts, causing multi-organ infections and sepsis. These infections are particularly difficult to treat when caused by multi-drug resistant strains, which are common as Kp is able to acquire resistance to most antibiotic classes, including extended spectrum beta-lactams and carbapenems. Several distinct multidrug-resistant Kp clones have been isolated in hospitals worldwide, making this bacterium a major public healthcare burden. Kp strains harboring blaKPC, a plasmid gene encoding a carbapenemase, pose a particularly high risk to public health, and recently a Kp strain that is resistant to all 26 antibiotics licensed in the US has been isolated [13, 14]. The most widespread KPC producers are isolates from clonal group 258 (CG258), which is known to have spread throughout the Americas, Europe, Asia and elsewhere [15–18]. CG258 belongs to KpI, and includes isolates belonging to Sequence Type 258 (ST258), ST11, ST512 and ST340, all of which have been documented to cause hospital outbreaks [19].

Whole genome sequencing has revealed that CG258 has experienced four large recombination events, each spanning at least 100Kb and up to 1.5 Mb [3–5, 7]. ST258 emerged after a >1Mb recombination between an ST11-like recipient, and an ST442-like donor [4]. However, it is currently unclear to what extent these large recombination events are actually associated with the epidemiological success of the clinically important CG258 clones [6]. More broadly, questions remain concerning the patterns of gene flow within the Kp population, and whether certain lineages are more or less likely to act as either donors or recipients. Although the data is currently limited, Holt and colleagues [9] observed that large recombinations tend to be less common between unrelated Kp phylogroups, and argued that gene flow in the Kp population is likely to be structured by biological and/or ecological barriers [9]. In order to investigate the large recombination phenomenon in Kp, we analysed over 800 genomes from this species, focusing on recombination analysis, donor identification, gene presence/absence. We show that large recombination events in Kp CG258 led to the emergence of several epidemiologically relevant lineages and provide evidence that suggests that donor gene composition may affect the successfulness of the hybrid strains.

## Results

### The sample analysed

The genomes of 12 ST11 isolates and one ST442 isolate, collected from France, Brazil, China and Japan between 1997 and 2014, were sequenced and assembled (Table S2). These 13 novel genomes were added to a collection of 821 genomes, to represent the genomic diversity of Kp (Table S3). A SNP-based phylogeny of this sample was consistent with previous studies ([9]; Figure S1). The vast majority of the isolates corresponded to a large radial expansion within KpI, with KpII and KpIII clearly distinct at the end of long branches.

Although the 834-genome dataset represents the diversity within the publicly available datasets, there exists a significant bias towards sampling clinical isolates. In order to control for this, we sub-sampled 394 Kp representative genomes based on pairwise SNP distances, such that no two genomes sharing fewer than 5 SNP differences were kept (see Methods). A subset of 60 CG258 strains was then extracted from the 394-genome dataset. Core SNPs were called independently for the global (394 strains) and CG258 (60 strains) datasets, and these SNPs were used for phylogenetic reconstruction using Maximum Likelihood (Figures 1 and 2).

**Figure 1.**
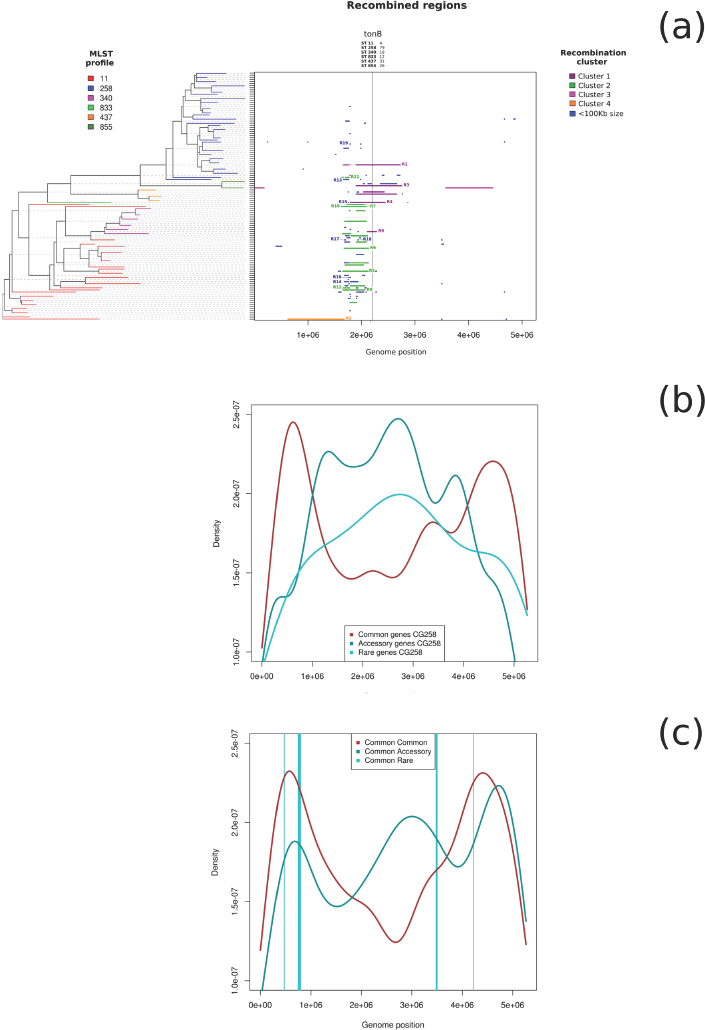
Recombination analysis, performed with ClonalFrameML. On the left, the phylogenetic tree obtained with RaxML with branches coloured on the basis of the MLST profiles. On the right, the plot of the recombined regions, coloured on the basis of the involved genomic regions as grouped by PVClust (see materials and method).

**Figure 2.**
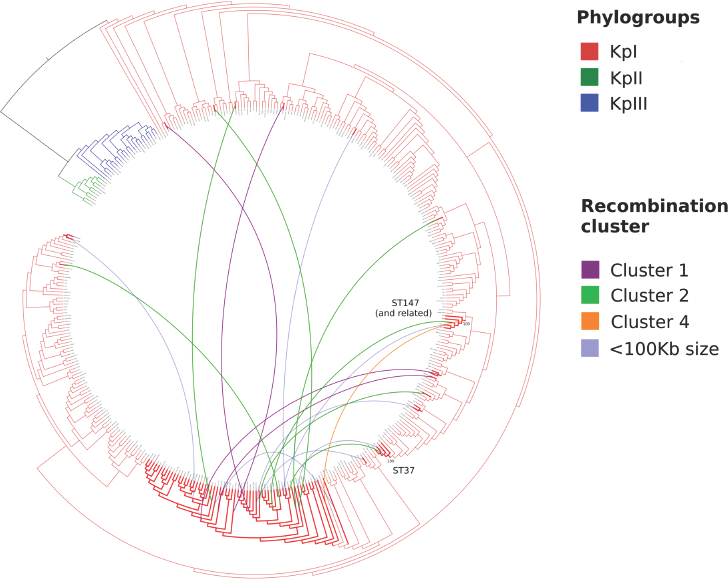
RaxML tree of 394 *Klebsiella pneumonaie* strains, selected from the global genome database to be representative of the genetic variability of the species. The branches of the tree are coloured on the basis of the phylogroup: red for KpI, green for KpII and blue for KpIII. The CG258 clade, on the bottom, is highlighted in red. The edges connect donors and recipients as identified in this work. The edge colour corresponds to the recombination cluster (see material and methods).

### Patterns of recombination within CG258

Recombination was detected within the 60 CG258 genomes using ClonalFrameML [20] (see Methods). A total of 119 recombination events were detected, 65 on internal nodes and 54 on terminal branches of the tree (Figure 1). These recombination events encompassed 63% (n = 3,344,516) of the 5,263,229 sites in the core genome alignment. In particular, 2,195,715 positions resulted recombined only in one branch of the tree, while 391,724 in at least 10 branches. Thirty of the 119 (25%) recombinations were sized > 100Kb (~2% of the reference genome) and, among them, six have >500 Kb size (~10% of the reference genome length) and two > 1 Mb (~20% of the genome). Thus, these initial observations confirmed that large recombination events have occurred relatively frequently within CG258.

Figure 1 clearly illustrates that the recombination events are not randomly distributed across the genome but are highly clustered. Ninety-five of the 119 (80%) total predicted recombination events occurred in a 1,185,000 sized region (23% of the genome), between positions 1,575,000 and 2,760,000 of the reference genome, and the same region contains 27 of the 30 large recombinations (>100 Kb) localized between the positions 1,660,631 and 2,750,819. In fact, bootstrap-based hierarchical clustering analysis reveals two highly supported clusters corresponding to two partially overlapped genomic regions, as shown by green and purple lines in Figure 1. Cluster 1 is composed of 7 recombination events spanning the region from 1,660,631 to 2,750,819 (1,090,188 bp), while cluster 2 includes 20 recombination events from 1,629,115 to 2,131,208 (502,093 bp). Thus, we divided the highly recombined region in two sub-regions: the “overlapped Cluster1-2” subregion, from 1,629,115 to 2,131,208, involving both “Cluster1” and “Cluster2” recombinations, and the “Cluster1 only”, from 2,131,209 to 2,750,819, involving only “Cluster1” recombinations.

Our analysis on the 60 representative genomes of CG258 revealed that ST340, ST437, ST833 and ST855 have emerged from a ST11-like ancestor following the acquisition of genomic regions of at least 100 Kb. All these STs are single-locus variants of ST11, the single discrepant allele being at tonB, which is located within the recombined region (Figure 1). Furthermore, three basal branches of the CG258 tree, corresponding to US-MD-2006, JM45 and CHS_24 strains, resulted completely free of large-recombination signals (Figure 1), suggesting that genomic features shared among them, such as gene content, were probably also common to the ST11 ancestor genome (for this reason we will refer to these three strains as “ST11-ancestor-like”).

### Identifying the origins of imported DNA in CG258

In order to identify the donors of the recombination events detected within CG258, we used a phylogenetic approach based on the core SNPs within each of the recombined regions. Using this method, we were able to robustly identify the donors of 19 recombination events, which we will refer to as “Recombination with Identified Donor” or RID (see Table 1, Table S1 and Figure S3 for information about RIDs donors and recipients, and Figure S4-S22 for trees). RID1 and RID19 correspond to previously identified donor / recipient pairs, thus confirming the soundness of our approach [3, 4].

In order to visualize the flow of large recombinations towards CG258, we plotted the global tree of Kp (including all the strains used for recombination analysis or donor identification) and connected donors and recipients with links (Figure 2). We found that only two highly supported lineages, ST147 and ST37, resulted involved in multiple recombination events.

**Table 1.**
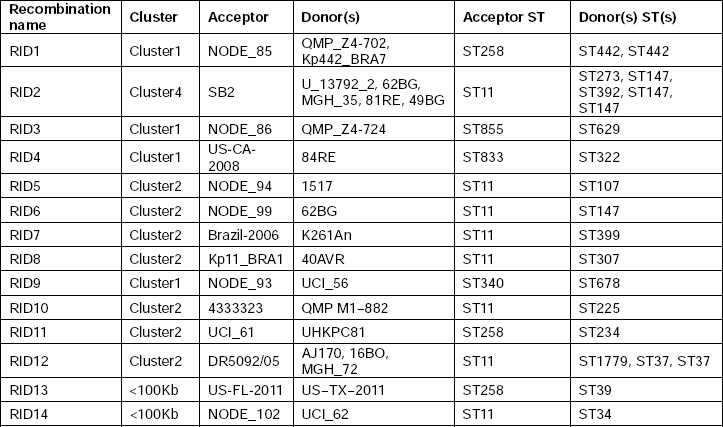
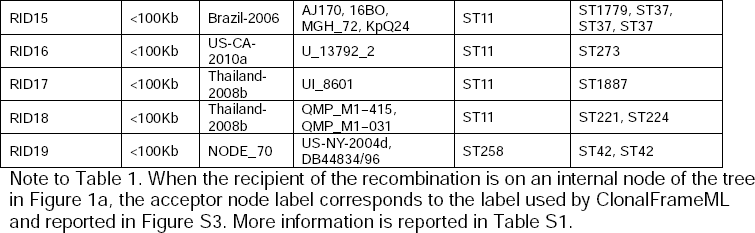
Main information about the recombinations for which the donors were identified (RIDs).

### Gene presence/absence analysis

We investigated whether divergent gene compositions between donors and recipients can represent a genetic barrier for large recombinations between non-CG258 (possible donors) and CG258 strains (possible recipients). We compared the genomic localizations of large recombinations and the positions of the genes commonly present among CG258 but less frequent among non-CG258 strains. More specifically, the 834 genomes included in the global data set were subjected to orthologue analysis, and genes were classified as “common” (present in >= 95% of the strains), “accessory” (< 95% and >= 5%) and “rare” (<5%), considering separately the CG258 and non-CG258 strains (classification of genes as “common” and “accessory” is according to Holt et al. 2015). Subsequently, the CG258 “common” genes were divided into “common-common” if classified as “common” also among non-CG258 strains, as “common-accessory” if classified as “accessory” in non-CG258, and as “common-rare” if “rare” in non-CG258 strains. Considering the observed high frequency of recombination, we expected that gene co-presence among recombination donors could introduce biases in the classification of CG258 “common” genes. Thus, we considered only CG258 “common” genes likely present into the ST11 ancestor, i.e. those shared among the three ST11-ancestor-like strains described above.

Within CG258, a total of 2453 common, 6786 accessory and 22539 rare genes were identified. In order to study the genomic localization of these gene categories, we considered only the genes present on the reference genome (2404/2453 common, 2738/6786 accessory and 61/22523 rare genes). The common, accessory and rare genes showed an evidently uneven distribution along the genome: common genes clustered around the origin of replication (ORI), while accessory and rare genes are more frequent in the central part of the genome (Figure 1b).

Among the 2453 genes classified as common in CG258, 2385 resulted present in all the ST11-ancestor-like strains and 1327 were classified as “common-common”, 977 as “common-accessory” and 81 as “common-rare”. The genomic positions of these genes present in the reference genome (1309/1397 common-common genes, 948/977 common-accessory and 80/81 common-rare genes) were then retrieved and compared to the positions of large recombinations. The distributions of common-common, common-accessory and common-rare genes show an interesting pattern (Figure 1c), in particular within the highly recombined genomic region described above (see “Patterns of recombination within CG258” paragraph). Indeed, in correspondence of the less frequently recombined genomic region called “Cluster 1 only”, common-common genes frequency reaches its minimum and common-accessory genes frequency show a local maximum (Figure 1c). Furthermore, no common-rare genes are localized within the highly recombined region (Figure 1c).

Out of the 948 common-accessory genes, 183 are localized within the highly recombined region. Non-CG258 strains show a variable pattern of presence of these genes (Figure S23), and statistical analyses revealed that: (a) strains involved as donors in multiple recombinations (ST147 and ST37) harbored significantly more of these genes that the other donor strains (Wilcoxon test, p-value < 0.05, boxplot in Figure S25); (b) within the highly recombined region, genes localized within the less frequently recombined “Cluster 1 only” sub-region, resulted harbored by significantly fewer non-CG258 strains in comparison to those localized within the more frequently recombined “overlapped Cluster1-2” sub-region (Wilcoxon test, p-value < 0.01 – boxplot in Figure S26). On the other hand, the common-rare genes heatmap (Figure S24) shows that rare genes are particularly frequent in some non-CG258 lineages, but an evident pattern with donors is not detectable.

Common-accessory and common-rare genes present in the reference genome were then annotated against Clusters of Orthologous Groups (COG) and pie charts of COG pathways abundances were plotted (Figure S27 and Figure S28, respectively). Finally, COG pathways abundances of common-accessory genes localized inside and outside the highly recombined region were compared and no significant difference was found (Chi-square test, p-value > 0.05).

## Discussion

Whole genome sequencing has revealed an unprecedented degree of genome plasticity in Kp, both in terms of the rates of horizontal gene transfer affecting the pan-genome, and in terms of the rates of homologous recombination in the core genome. Most strikingly, this species undergoes very large recombination events, affecting up to 20% of the genome [3–5, 7, 9]. However, it remains unclear to what degree these large recombination events are responsible for the epidemiological success of lineages such as CG258, nor whether gene flow is in some way structured within the broader Kp population.

To investigate if gene flow affects large recombinations, we subjected a representative subset of 60 CG258 genomes (selected from more than 400 CG258 genomes, sequenced as part of this study or retrieved from publicly available databases) to recombination analysis, donor identification and gene presence/absence analysis, obtaining an overview of the large recombination phenomenon in the lineage, and thus novel epidemiological and the evolutionary insights.

Our analyses reveal that large recombination events (>100 Kb) occur commonly in Kp, strongly highlighting them as a persistent mechanism of diversification in the CG258 clade. Furthermore, we found that, in addition to ST258, four other Kp lineages of epidemiological relevance (ST340, ST437, ST833 and ST855) emerged by large recombination events, suggesting that large recombinations have an important role in generating epidemiologically relevant clones.

Based on the results obtained from recombination analysis, donor identification and gene presence/absence analysis, we propose the hypothesis that a different gene composition between donor and recipient can limit the success of the emerged hybrid strains. According to this hypothesis, the donor of a successful large recombination must possess, within the exchanged genomic portion, the genes necessary for the survival of the recipient strain (genes already present within the replaced portion of the recipient genome). Following this model, the number of successful donor-recipient combinations is likely limited, as well as the number of possible emerging hybrid strains. We graphically illustrate the proposed model in Figure 3.

**Figure 3.**
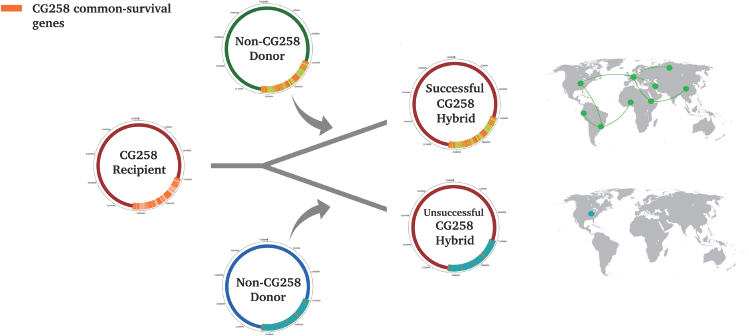
Graphical representation of the hypothesis proposed in this work: after a large recombination, the success of the emerged hybrid strain depends on how many survival genes of the recipient are present within the genomic region provided by the donor. The recipient strain genome is represented as a red circle (on the left), the genomic region that is replaced is in light red and the survival genes within this region are represented as orange lines. Two possible events are represented: (a) a large recombination involving a donor (in green) harbouring many survival genes produces a successful hybrid strain able to spread worldwide; (b) a large recombination involving a donor (in blue) harbouring a few survival genes produces an unsuccessful hybrid strain.

The rationale for this hypothesis can be summarized as follow: (a) the acquisition of a large genomic region can produce changes in the gene content into the recipient genome [2] (Figure 1b) and it is reasonable to assume that such phenomenon would be particularly prominent in bacterial species with highly variable gene content, such as Kp [9]; (b) gene content changes can dramatically affect the fitness of the recipient strain, thus we can assume that the large recombinations observed are just the successful ones, because a recombination that causes the loss of genes fundamental to the recipient (“putative recipient survival genes”) would cause a significant reduction of the fitness of the emerged strain; (c) the absence, in the transferred genome region, of genes required for recipient survival could represent a genetic barrier to large recombinations.

The evidences that led us to formulate, and then to support, this hypothesis are discussed below.

1. Discontinuity in recombination frequency along the genome suggests the existence of a genetic barrier.

Large recombinations involve more frequently a specific 1,5 Mb sized genomic region (here called “highly recombined region”, Figure 1a). Within this region, clustering analysis revealed the existence of two partially overlapping, but well-delimited, sub-regions presenting different rates of recombination (Figure 1a) (on the basis of the clustering result we called the most frequently recombined ones “overlapped Cluster1-2”, and the other one “Cluster1 only”). Despite the higher frequency of recombination of the entire region can be explained by a strong diversifying selection due to the presence of the capsule genes [7], the discontinuity between the two sub-regions (both involving the capsule genes), needs an additional explanation. Indeed, it is reasonable that, in a bacterium able to exchange large genomic portions, the existence of a genomic locus with a high rate of successful recombination (such as the locus of the capsule genes) can produce an hitchhiking-like effect of adjacent loci: the probability that a locus is involved by a successful large recombination event is expected to decrease with the distance from that high frequently exchanged locus. The existence of two well-delimited genomic sub-regions with different recombination frequencies suggests a different mechanism, such as the existence of a genetic barrier to large recombinations between these two regions.

2. The absence of a pattern between exchanged regions and donor lineages suggests that the genetic barrier could be localized on the recipient genome.

We robustly identified the donors of 19 recombination events in the CG258 lineage (Figure 2). We found that the lineages ST147 and ST37 are donors in multiple recombination events. No association was observed between the localization of the large recombinations and the phylogenetic position or ecological origin of the donors. This result supports the idea that the localization of large recombinations is affected by genomic factors of the recipient genome.

3. Testing the hypothesis: is there a relationship between gene content and recombination frequency?

In order to test our hypothesis, we focused on CG258 “putative surviving genes”, identified as the genes present in > 95% of all the CG258 strains included in the study (>400 strains). In particular, we investigated their localization within the “highly recombined region” and their frequency among the non-CG258 strains, possible donors of large recombinations. In order to minimize possible biases due to gene content variations caused by large recombination, we decided to include in the analyses only the CG258 “putative surviving genes” harbored by the three deep-branching CG258 strains, for which recombination analysis did not report evidence of large recombination in their evolutionary history (see Results). Indeed, we assume that genes present in the genome of these three strains were also likely present in the unrecombined GC258 ancestor. We will refer to these genes as “CG258 common genes”.

We found that many of the CG258 common genes are localized around the origin of replication, outside the “highly recombined region” (Figure 1b). This distribution can be explained in two ways: a) the high number of recombinations increased the gene content variability of the “highly recombined region”; b) the lesser content of survival genes within this region makes it more replaceable.

In order to discriminate between these two possible explanations, we sub-classified the CG258 common genes on the basis of their frequencies among non-CG258 strains (we classified the CG258 common genes as “common-common” if present in > 95% of the non-CG258 strains, as “common-accessory” if > 5% − <= 95% and as “common-rare” if < 5%). Indeed, if the successful rate of a large recombination is affected by donor gene content within the exchanged genomic region, we should aspect to observe a pattern between the recombination frequency of a CG258 genomic region, and the frequency among possible donors (non-CG258 strains) of the genes localized within that genomic region.

We found a pattern between the genomic localizations of large recombinations and those of common-common, common-accessory and common-rare genes. Indeed, (a) the absence of common-rare genes within the highly recombined region suggests that the presence of these genes could be a strong barrier for large recombination events in a CG258 recipient; (b) the higher frequency of common-accessory genes and lower frequency of common-common genes within the lesser frequently recombined “Cluster1 only” genomic region, suggests that only a limited number of possible donors could exist for this genomic region.

Two additional lines of evidence support the hypothesis of gene content as a genetic barrier: (a) common-accessory genes localized within the “Cluster 1 only” genomic region are less frequently present in possible donors (non-CG258 strains), in comparison to those localized into the highly recombined “overlapped Cluster1-2” region; (b) the strains of the donor lineages involved in multiple large recombinations (ST147 and ST37) harbor more common-accessory genes localized within the highly recombined region than the other donors.

Common-accessory genes localized within the highly recombined region resulted to belong to multiple fundamental pathways, highlighting how a large recombination can affect several compartments of the recipient metabolism.

## Conclusions

Large recombinations frequently occurred during the evolution of *K. pneumoniae* clonal group 258, leading to the emergence of novel lineages. Our work reveals that large recombinations occurred with higher frequency in specific Kp lineage pairs, and that the prevalence of these events is not evenly distributed across the Kp genome. Here we propose that this pattern could be explained if we consider the different gene content between recipients and donors as a barrier for large recombinations. This first reconstruction of a network of large recombination events in Kp provides a novel point of view on this phenomenon, highlighting the importance of such an approach for investigating the evolution of recombinogenic bacterial species.

## Material and Methods

### Genome sequencing

Thirteen Kp hospital isolates were obtained from Brazil (eight isolates), China (three isolates), Japan (one isolate), and France (one isolate), based on their MLST profile: twelve ST11 and one ST442. DNA was extracted using a QIAamp DNA mini-kit (Qiagen) following the manufacturer’s instructions. Whole genomic DNA was sequenced using an Illumina Miseq platform with a 2 by 250 paired-end run after Nextera XT paired-end library preparation. Paired-end genomic reads were assembled using MIRA 4.0 software [21].

### Reconstruction of Kp species and CG258 representative databases

663 genome assemblies and 158 genome reads datasets were retrieved from the NCBI and Patric databases, for a total of 821 strains collecting of the entire known Kp species genomics variability at May 2015 (i.e. including strains belonging to the KpI, KpII and KpIII phylogroups). The downloaded reads were assembled using Velvet software [22]. All genomes were then merged in a 834 genomes dataset and aligned against the complete genome of the Kp reference strain 30660/NJST258_1 [3] using progressiveMauve [23]. The multi-genome alignment and the core Single Nucleotide Polymorphisms (core SNPs) alignment were obtained using the in-house pipeline described by Gaiarsa and colleagues [5]. The core SNP alignment was then subjected to phylogenetic analysis using RAxML version 8.0.0 [24] using the ASC_GTRGAMMA model and 100 bootstrap replicates.

In order to remove oversampled clones and clades and to obtain a dataset of manageble size while maintaining the information of the entire genomic variability of the species, a Kp species representative genome database (from now on referred to as species database) was constructed using the following procedure: (a) SNPs distance matrix among the strains was obtained using the R library Ape [25, 26]; (b) a recursive process of strain selection was performed, removing strains with less then five SNPs distance from others. The strains belonging to CG258 were then manually extracted from the species database, thus obtaining a representative selection of CG258 strains (from now “CG258 database”).

### Recombination analysis

The CG258 database and two outgroup strains (18PV and K102An) were subjected to reference-based genome alignment, SNP calling and phylogenetic analysis, as above. The obtained tree was rooted on the outgroups and the outgroups were then removed to obtain a representative CG258 tree. Recombination analysis was performed using the ClonalFrameML software [20], setting the transition/transversion rate as calculated by PhyML [27]. The positions of the recombinations of more than 100 Kbps were retrieved, compared and subjected to clustering analysis: the start and end positions of the recombinations were used to compute distance matrix using the Manhattan distance, and hierarchical clustering (with p-values) algorithm implemented into the Pvclust [28] function was used to group the recombinations. Highly supported clusters were identified setting the approximately unbiased index threshold at 0.99. The analyses were performed using R [26].

### Identification of the donors of the recombinations

In order to identify the Kp donors of the recombinations we performed an ad-hoc phylogenetic analysis. The genomes of the species database were aligned to the reference genome and core SNPs were called, subjected to ML phylogeny using FastTree software [29] with 100 bootstrap (we will refer to the obtained tree as “species database tree”). For each recombination, the core SNPs called within the recombined region were extracted and subjected to phylogenetic analysis, using FastTree software [29] with 100 bootstrap. Each resulting tree was manually analysed as follow: (a) the CG258 recipient(s) of the recombination was (were) identified on the tree; (b) when a highly supported (> 75 bootstrap) monophylum including all the recipients and one or more non-CG258 strains was detected, the non-CG258 strains were considered as donors of the recombination.

Recombinations identified on the same branch of the CG258 tree, localized on adjacent genomic regions, and sharing the same donors, were merged, as likely originated from a single recombination event. For these recombinations, the donor identification procedure was repeated considering a novel recombined region, ranging from the beginning of the first recombination to the end of the second one.

The reliability of the identified donors was assessed testing if they make a monophyletic clade on the smaller global tree.

### Gene presence absence analysis

The 834 genomes included into the global genome database were subjected to Open Reading Frame (ORF) calling using Prodigal [30], and then to ortholog analysis using Roary software [31] after the annotation with PROKKA [32]. The obtained gene presence/absence matrix was then analysed as follow. We split the matrix into two sub-matrices: the first one including CG258 strains and other one non-CG258 ones. From each sub-matrix we classified the genes as “common” (present in >=95% of the strains), “accessory” (<95% and >=5%) and “rare” (<=5%). The positions of the CG258 “common”, “accessory” and “rare” genes present on the reference genome were retrieved merging the information from the Roary output and the reference annotation file, using an in-house Perl script. The cumulative distribution of the positions of the genes of each category along the reference genome was obtained and plotted using R [26].

Then we classified the CG258 common genes as follow: “common-common” if classified as “common” among non-CG258 strains, “common-accessory” if classified as “accessory” and “common-rare” if “rare”. Then, the positions of these genes of the reference genome were retrieved as described above, and gene categories distributions and genes occurrence among non-CG258 strains were plotted using R [26]. Furthermore, we compared gene composition of Kp lineages using Wilcoxon and Chi Square tests using in R [26].

## Acknowledgments

The work was supported by the Romeo ed Enrica Invernizzi Foundation.

## Captions to supplementary figures

**Figure S1.** Maxumum Likelihood tree of 834 Kp strains obtained with RaxML software.

**Figure S2.** Result of the bootstrapped clustering performed on the >100Kb sized recombination. The recombinations were clustered on the basis of the start and end positions on the genome. The clustering analysis was performed using PVClust R package, and the clusters were identified setting the au threshold at 0.99. The two major clusters, Cluster 1 and Cluster 2, are highlighted in violet and green respectively.

**Figure S3.** Maximum Likelihood tree (without branch length information) obtained from RaxML and subjected to recombination analysis with ClonalFrameML. The node labels correspond to those gave by ClonalFrameML to identify the acceptors of the recombinations, and correspond to acceptor names used in Table 1.

**Figure S4-S22.** (a) Maximum Likelihood phylogenetic tree of the Kp strains included in the donor identification analysis, performed using the core SNPs called within the recombined genomic region. (b) Blow-up of the clade of the phylogenetic tree (reported in figure a) that contains the recipient(s) of the recombination (in blue) and the putative donor(s) (in red) of the recombination.

**Figure S23.** On the left, SNP-based Maximum Likelihood phylogenetic tree, obtained using RAxML software, including all the non-CG258 strains from the global Kp strains dataset. In the middle, heatmap plot of the common-accessory genes presence/absence among the strains. Colors identified recombination donors: green, strain not donor; violet: donor of a large recombination (>100Kb size) into “Cluster 1” genomic region; green, “Cluster 2”; orange, “Cluster 4”; blue, donor of recombination <100Kb size; red, donor that belong to a lineage involved in multiple recombination event. On the right the number of genes harboured by each strain is reported.

**Figure S24.** On the left, SNP-based Maximum Likelihood phylogenetic tree, obtained using RAxML software, including all the non-CG258 strains from the global Kp strains dataset. In the middle, heatmap plot of the common-rare genes presence/absence among the strains. Colors identified recombination donors: green, strain not donor; violet: donor of a large recombination (>100Kb size) into “Cluster 1” genomic region; green, “Cluster 2”; orange, “Cluster 4”; blue, donor of recombination <100Kb size; red, donor that belong to a lineage involved in multiple recombination event.

**Figure S25.** Boxplot of the amount of common-accessory genes among donor strains, dividing those involved in multiple recombination events from the others. Wilcoxon test, p-value <0.01.

**Figure S26.** Boxplot of the mean amount of non-CG258 strains harbouring common-accessory genes localized within the “Overlap Cluster 1 and 2” and within the “Only Cluster 1” subregions. Wilcoxon test p-value < 0.05.

**Figure S27.** Pie chart of the Clusters of Orthologous Groups (COG) pathways of the 183 common-accessory genes localized within the highly recombined region.

**Figure S28.** Pie chart of the Clusters of Orthologous Groups (COG) pathways of the 80 common-rare genes present into the reference genome.

